# Leveraging the Elastic Deformability of Polydimethylsiloxane Microfluidic Channels for Efficient Intracellular Delivery

**DOI:** 10.1101/2022.07.22.501113

**Authors:** Hashim Alhmoud, Mohammed Alkhaled, Batuhan E. Kaynak, M. Selim Hanay

## Abstract

With the rapid development of microfluidic based cell therapeutics systems, the need arises for compact, modular, and microfluidics-compatible intracellular delivery platforms with small footprints and minimal operational requirements. Physical deformation of cells passing through a constriction in a microfluidic channel has been shown to create transient membrane perturbations that allow passive diffusion of materials from the outside to the interior of the cell. This mechanical approach to intracellular delivery is simple to implement and fits the criteria outlined above. However, available microfluidic platforms that operate through this mechanism are traditionally constructed from rigid channels with fixed dimensions that suffer from irreversible clogging and incompatibility with larger size distributions of cells. Here we report a flexible and elastically deformable microfluidic channel, and we leverage this elasticity to dynamically generate temporary constrictions with any given size within the channel width parameters. Additionally, clogging is prevented by increasing the size of the constriction to allow clogs to pass. By tuning the size of the constriction appropriately, we show the successful delivery of GFP-coding plasmids to the interior of three mammalian cell lines and fluorescent gold nanoparticles to HEK293 FT cells all while maintaining a high cell viability rate. This development will no doubt lead to miniaturized intracellular delivery microfluidic components that can be easily integrated into larger lab-on-a-chip systems in future cell modification devices.

## Introduction

Intracellular delivery of biologically active cargo is essential for gene therapy and tissue engineering as well as to study fundamental cellular function. The cellular membrane regulates and blocks the free transit of exogenous materials into the cytosol. In order to transport cargo across the cellular membrane without negatively impacting cellular function, there has been extensive research aimed at developing non-invasive cargo delivery methods. These methods can be broadly classified as either biochemically based, or physically based. Biochemical methods take advantage of existing cell machinery or functions to force the internalization of cargo and typically includes the use of lipid/polymer nanocarriers, viral vectors and pore-forming agents such as detergents and ligand conjugates.^[1-4]^ Biochemical approaches suffer from their fair share of issues such as slow delivery, complexity in synthesis, limitation in cargo types, and inconsistent results, in addition to eliciting adverse immune responses from the host in the case of viral vectors.^[3, 5]^

Alternatively, physical approaches rely on causing transient cell membrane openings either through physical penetration (such as the case with nanoneedles or microinjections)^[6]^, or through membrane permeabilization. Electroporation,^[7-8]^ sonoporation^[9]^, and optoporation^[10]^ are some of the more well-known technologies by which membrane disruptions and permeations are induced temporarily. These permeations allow transmembrane transportation of a variety of cargos such as proteins, DNA and RNA, small molecules, and organic and inorganic nanoparticles. Physical approaches afford more flexibility in the choice of target cargo as compared to biochemical approaches, and typically tend to be simpler to implement.^[11]^ However, the above-mentioned physical approaches have also been linked to diminished cell viability rates since some of the membrane disruptions become irreversible. This is especially the case with electroporation^[12-13]^ and sonoporation^[14]^ due to irreversible membrane disruption. Another physical method: rapid cell deformation through constrictions, more commonly known as cell “squeezing” has been shown to create transient membrane pores that both allow cross-membrane transport at an efficient rate while also maintaining high cell viability.

The first report by Sharei *et al*.^[15]^ described successful intracellular delivery achieved via rapid cell deformation through a microchannel constriction. This was promptly followed by a multitude of studies on the various constriction geometries and their effect on cell survivability and intracellular transport efficiency while demonstrating the successful delivery of many types of cargo to a variety of cell lines^[16-20]^.

The delivery efficiency of cell squeezing platforms depended primarily on constriction size relative to cell diameters and the flowrate. It was shown that a constriction 30-70% of the target cell size was necessary for any significant permeabilization effects to take place^[18, 21]^. On the other hand, if the constriction was too small, cell viability plummeted. This meant that control over constriction size was paramount in such systems. However, the cell squeezing platforms reported so far rely on fixed channel geometries etched into a rigid material such as silicon or glass which meant that the channel geometry was suitable for only cells with certain size distributions. Additionally, a fine balance between flow rates and constriction geometries is required to maintain optimal cell permeabilization. Clogging in such channels has necessitated the use of very high flowrates and multiplexed parallel and complex channel designs. The high flowrates markedly reduced cell viability while the channel design rigidity made it vital to fabricate fine-tuned geometries for each cell type individually. These hinderances limited the devices’ versatility across multiple cell species and prevented channel reusability. Here, in order to resolve the issue of fixed channel geometries we leveraged the elastic deformability of polydimethylsiloxane (PDMS) channels to create constrictions with tunable sizes dynamically by applying mechanical force on the outer boundaries of the microfluidic device. This is accomplished by using simple macro sized linear actuators. By tuning and focusing the mechanical force onto a specific region of the channel (referred to as the compression region), the channel deforms inwardly with micrometer-scale precision to form a constriction in real-time while cells are flowing through the channel. We apply this mechanism to transfect three cell lines with a green fluorescent protein (GFP) coding plasmid and measure the transfection efficiency in light of the efficiency of a commercial transfection reagent. We also use the same mechanism to transport fluorescently tagged gold nanoparticles (AuNPs) to HEK293 FT cells, which is not achievable using the commercial transfection reagent.

## Results and Discussion

### Device design and operation

The elastic deformability of PDMS is perhaps one of the most characteristic properties of the material, which allows for many useful implementations in field of microfluidics.

In contrast to PDMS-on-glass microfluidic devices, PDMS-on-PDMS devices are entirely elastic, with the strength of the PDMS-PDMS bonding ensuring that the device can withstand high effective fluid flowrates without developing leaks. Additionally, the surface chemistry of PDMS is quite versatile and can be functionalized with ease using silane chemistry. These advantages, combined with PDMS’s inherent biocompatibility makes the material ideal for constructing fully flexible microfluidic devices for cell manipulation.

In this proof-of-concept, we set out to demonstrate that by carefully applying an appropriate amount of external mechanical force on a PDMS-on-PDMS microfluidic channel, the corresponding elastic deformation generates a temporary constriction at the point of mechanical pressure (compression region). This can be utilized to mechanically interact with cells passing through the described constriction. The overall concept is demonstrated in Figure 1a, where a straight microfluidic channel surrounded by force applicators deforms in shape upon being exposed to the compressive force. Furthermore, the inclusion of micro-features within the compression region to guide the type of interaction that is to take place, e.g. cell elongation.

**Figure 1:**
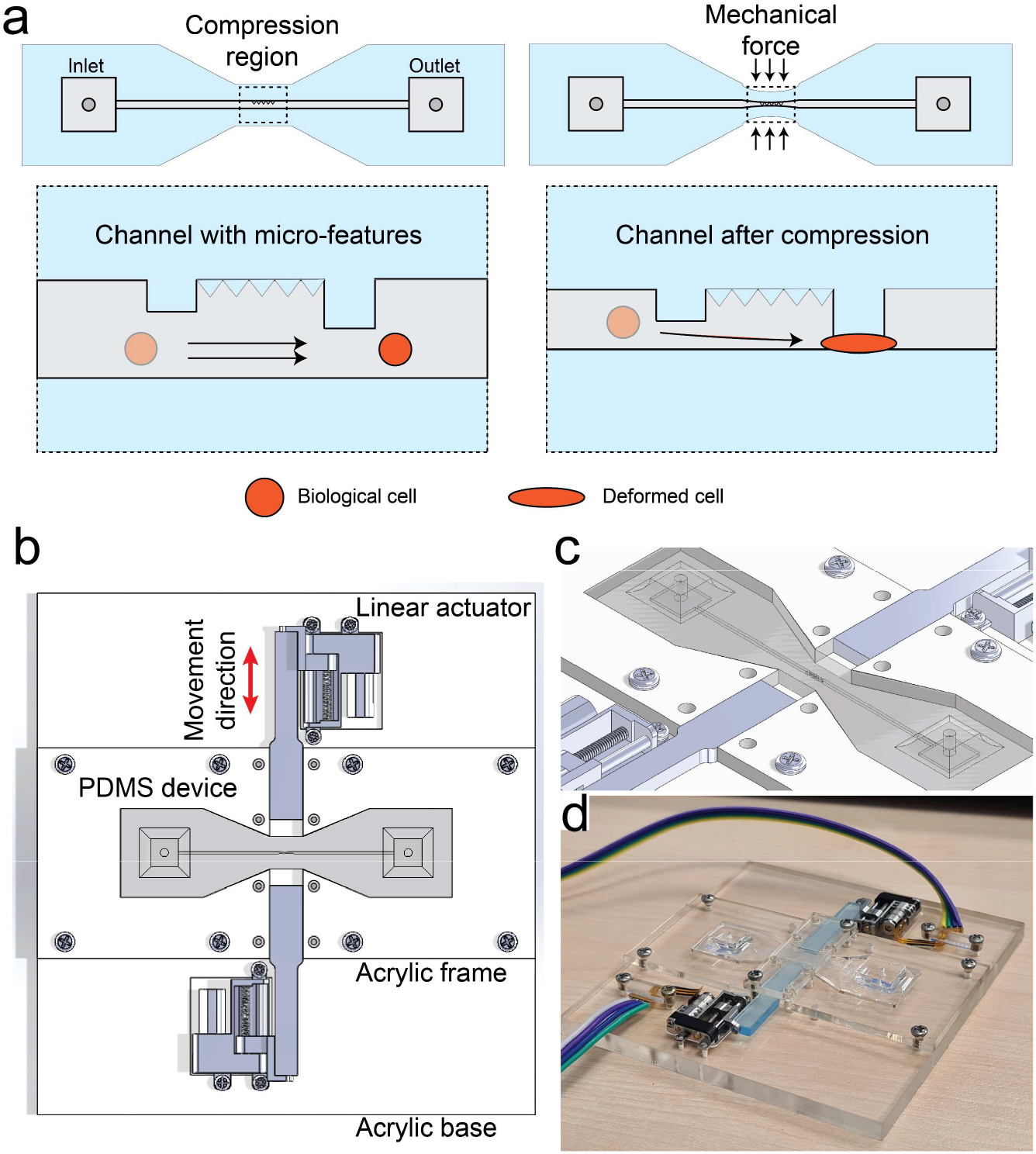
Schematic representing the PDMS device architecture in addition to the acrylic frame with the attached linear actuators. a) represents a top-view schematic of the PDMS device dog-bone shape before and after mechanical compression. The insets show a close-up of what is expected to occur inside the channel at the compression region as a result of applied mechanical force. The channel deforms inwardly to create a temporary constriction that forces cells to deform in turn to cross the constriction. b) is a schematic of the acrylic frame that was constructed to hold the PDMS device in place in addition to the linear actuators. c) represents an isometric view of the same assembly from (a) showing the depth of the 3d PDMS device with a thickness of ∼2 mm. d) is an optical photograph of the assembly showing the miniature linear actuators connected to the rigid mechanical arms and driven by external circuitry. As in the schematics from (b) and (c), the PDMS device is situated in the middle surrounded by an acrylic frame.

The overall device architecture is composed of two layers of PDMS bonded together, with the first containing an engraved 100 µm wide x 100 µm tall microfluidic channel that is 39 mm long (Figure S1a-e). The microfluidic channel terminates on both ends with square-shaped 5x5 mm reservoirs to simplify attachment to a fluid flow control system. The assembled device was cut into a dog-bone shape with the narrower region being aligned with the mid-point of the microfluidic channel and comprising the compression region.

Two miniaturized linear actuators (detailed in the Experimental section) are situated on opposite ends of the PDMS device and connected to rigid acrylic pusher arms that deliver compressive mechanical force against the compression region boundaries of the PDMS device along a single axis (Figure 1b-d). Any vertical movement as a result of PDMS compression is prevented by a clear acrylic rectangular piece secured in place by metal screws. The diameter of the constriction that forms as a result of mechanical compression is inversely related to the magnitude of the mechanical force being applied by the external linear actuators. The constriction size can then be described as a function of the Compression modulus (E) of the PDMS material which is defined by Hooke’s law as:

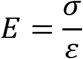

σ here represents the applied compressive stress in Pascals (Pa) and ε represents the strain (compression distance/original length). In a study by Wang *et al*.^*[22]*^ the modulus (E) was related to the base elastomer-to-curing agent weight ratio *n* in terms of MPa as:

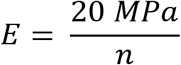

In order to optimize the PDMS elastomer mixing ratio so as to minimizing the force required to compress the channel, we performed finite element analysis on COMSOL software.

### Compression simulation

Using the analysis software, we applied displacement force ranging from 0 - 40 N and plotted the force as a function of channel width (0 – 100 µm) for elastomer mixtures ranging from 5:1 to 25:1 elastomer:cross-linker ratios. This is shown in Figure 2a. The simulation results showed that as the cross-linker concentration was increased, the elastic modulus (E) increased as expected, and the force required to completely collapse the channel increased. Lower cross-linker concentrations (15:1 to 25:1) showed force requirements plateauing at 10 ± 4 N. However, in practice, mixtures with cross-linker concentrations lower than 15:1 become extremely fragile. For this reason, the 15:1 ratio was the most robust and most compressible compromise to fabrication the device. The total force required to completely compress the 100 µm wide channel constructed from the 15:1 mixture was calculated to be ≈ 14 N. The two linear actuators attached to the device are each capable of delivering 10 N of force at maximum current draw. Combined, operating both actuators while modulating the step size allowed for very fine control over the resulting constriction size. Additionally, they allowed for the channel to remain within the field of view of a microscope objective while under compression due to symmetry.

**Figure 2:**
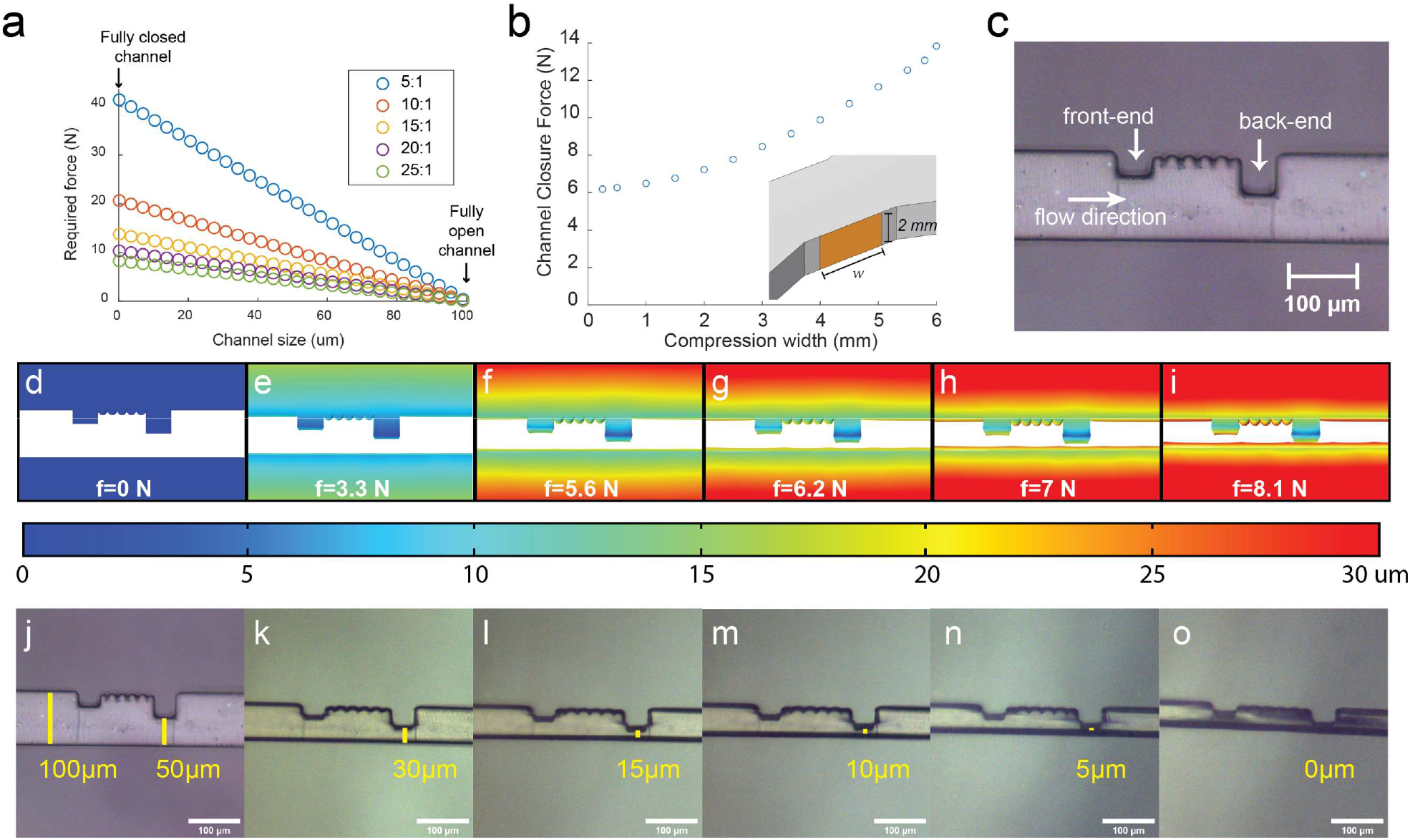
a) represents the linear relationship between the applied external mechanical force, and the width of the constriction forming within a 100 µm-wide channel as shown by the simulation at various PDMS mixing ratios. b) describes the force required to fully close the 100 µm channel with a 15:1 mixing ratio as a function of the width of the pusher arm tip in contact with the PDMS given that the tip height is fixed at 2 mm. c) an optical microscope image representing the typical configuration of the micro-features that facilitate mechanical interactions with cells passing through the region. d-i) is a rendered top-view of the channel that shows the simulated profile as more and more mechanical pressure is externally. The heatmap corresponds to the total displacement experienced by each point of the overall PDMS channel structure as a function of the applied force indicated at the bottom of each frame. The observed bulging is consistent with what is observed experimentally. j-k) are optical microscopy images showing the compression region as more external mechanical pressure is applied by the linear actuators. The channel within the compression region is constricted and the hydraulic diameter is reduced from 100 – 50 µm from wall to wall, while the distance between the largest square and the opposite wall drops consistently from 50 – 0 µm. The constriction size was purposely set to the desired values before taking the snapshot in each frame to demonstrate the level of control provided by this architecture to tune the constriction size.

Next, the contact area between the tip of the pusher arms and the PDMS device was optimized. The contact area is comprised of the thickness of the PDMS device (2 mm) multiplied by the width of the pusher arm tip. This could range from 1 – 6 mm laterally. The plot relating the force required for a 15:1 device to compress fully as a function of compression width (mm, since height is fixed) is shown in Figure 2b. The curve showed that the force required bottomed out at 1 mm tip width. However, in practice the sharp 2 × 1 mm pusher arm tip caused tears in the PDMS device at high compression forces. Therefore, a 2 × 2 mm or 4 mm^2^ contact area between the tip of the pusher arm and the PDMS device proved to be the most suitable. This resulted in a notable decrease in the force required for complete channel compression (from 14 to ∼ 7.5 N).

Micro-features were further included in the design of the microfluidic channel in order to create a smaller distance between the channel wall and the tip of the tallest micro-feature (50 µm) and to provide more precise control over mechanical interactions with cells in the form of right-angled traps. The micro-features were comprised of a 50 × 50 µm square at the back-end of the compression region which was the last structure to come into contact with incoming cells in order to act as the main constriction (Figure 2c). At the front-end of the structure, another 30 × 50 µm rectangle was placed to focus the flow of the incoming cells hydrodynamically, and to prevent cells from moving backwards out of the constriction area due to potential reverse flows caused by the change in channel geometry. The space between the two micro-features was decorated with spikes to reduce the planar contact area available for cell stiction and reduce the likelihood of forming clogs (Figure 2c).

A geometrical simulation of the shape of the constriction at different compression ratios was performed to visually observe the shape of the constriction region. This is shown in Figure 2d-i, where the heatmap corresponds the displacement experienced by each point of the channel structure as a function of the total applied mechanical force. In the simulation, we observed a bulging effect (Figure S2) of the channel sidewalls and the micro-features since both were fixed to the channel structure from the top and bottom. We also observed that the displacement occurred at a gradient, where the PDMS in contact with the pusher arms experienced the most displacement, while the channel sidewalls and the micro-features experienced the least amount. This demonstrated that large displacements externally translate in small displacements within the channel, which provided a high degree of precision in terms of channel deformation by tuning the torque of the external motors. This was further confirmed by physical experiments whereby the PDMS channel was compressed from 50 µm to 30, 15, 10, and 5 µm with precision as is shown in Figure 2j-o. We also observed the bulging effect shown as out-of-focus shadows in the channel in Figure 2n and o.

Both the simulation and the physical experiment demonstrated that the proposed device configuration along with the torque of linear actuators were able to give precise control over the constriction width for all subsequent cell deformation experiments.

### Clogging prevention

PDMS surfaces are notoriously sticky especially when they come in contact with biological materials. Non-specific protein adhesion to PDMS has been reported as a significant issue that hinders PDMS-based microfluidics^[23-24]^. Furthermore, constricted channels that are wide enough for one or two cells suffer from the high probability of clogging due to inherent physical limitations. This clogging issue was observed in early flow cytometry systems where cells were flown through narrow channels to optimize optical analysis. Channel clogging necessitated the utilization of sheath fluid to ameliorate this issue which is now standard practice in most flow cytometers. For mechanical constrictions however, the use of sheath fluid is not possible due to the obvious difficulties with cell-constriction interactions under sheath flow. In fact, this problem remains one of the most pressing for “squeeze” based intracellular delivery systems^[25]^. In a series of studies regarding this issue, it was shown that particles larger than one third (> 1:3) of the channel will invariably develop clogs even at surprisingly low volume concentrations^[26-28]^. This factor, coupled with the inherent large size distribution of cells in a given culture usually ensures that a device channel with a small diameter constrictions will be clogged after some usage^[25]^.

However, with this proposed device architecture, the ability to dynamically increase and decrease constriction size resolves this fundamental problem of clogging entirely since the constriction size can be increased when clogs develop. Furthermore, in order to solve the PDMS adhesion problem, the channels were chemically passivated with 1H, 1H, 2H, 2H-Perfluorooctyltriethoxysilane following the protocol described in the experimental section. The reaction scheme is shown in Figure S3.

After this procedure we observed a significant reduction in cell-channel adhesion or clumping even at the smallest constriction size (5 µm). In general, there was no observed adhesion or clumping behavior between the cells and at the cell/inner wall interface except in a few cases where one or two cells would adhere to the back-end micro-feature for a few seconds (as shown in the supplementary video) before being dislodged by fluid flow. The combination of passivated channel walls and the variable constriction size ensured that the PDMS device operated smoothly without developing clogs even at low flowrates (as low as 1 µL/min). Sample sizes as large as 5 mL with cell concentrations of 10^6^ cells/mL were easily treated with the device constriction at a rate of 1 mL/25 min using a maximum flowrate of 40 µL/min (Figure S4). An operator was necessary to control the constriction diameter in case of clogging, however this process will be eventually automated using image recognition software or through real-time flowrate measurements coupled to a feedback loop.

### HEK293 FT cell membrane permeation

The ability to elastically deform the compression region of the device on demand allows for an advanced level of control over individual cells that pass through the generated constriction. Using the device as intended enables the observation of changes that occur to cells as a result of morphological deformation in real time. To observe this process, we utilized a Human Embryonic Kidney (HEK293) cell line modified to be easily transferred to suspension through simple agitation without the use of Trypsin (designated as HEK293 FT). The cell line was suspended in Dulbecco’s Modified Eagle Medium (DMEM) supplemented with 20% fetal bovine serum (FBS) and containing 0.3 µg/mL of propidium iodine (PI) stain. PI stain is a DNA intercalating stain that exhibits peak fluorescence at 636 nm (red) upon intercalation. However, PI is a non-membrane permeable stain normally (i.e. cannot cross healthy cell membranes). Therefore, PI staining was an ideal indicator of membrane permeabilization. The suspended cells were introduced into the channel at a flowrate of 12 µL/min which provided a good balance between cell velocity and the ability to view cells under an optical microscope. A simplified schematic (Figure 3a-d) represents a compressed channel (10 µm constriction size) and shows the stages of morphological deformation that cells experience as observed under the optical microscope (Figure 3e-g). Cell deformation occurs rapidly as the cell encounters the corner of a square-shaped micro-feature located at the center of the compression region of the channel. Liquid flow forces the cell to traverse further through the constriction where its morphology begins to flatten to match the clearance between the tip of the micro-feature and the opposite inner wall of the channel. This rapid deformation has been previously shown to result in the creation of transient pores through the cellular membrane^[15, 19-21, 29]^. Passive diffusion of the desired cargo occurs rapidly at this stage from the extracellular media surrounding the cell along the concentration gradient. The results are shown in Figure 3h and the supplementary video where cells that were driven through the 10 µm wide constriction immediately began to fluoresce in the red channel as they exited the constriction. This confirmed that indeed the PI stain was entering the cell cytosol and interacting with cellular DNA as a result of the cell deformation through the constriction.

**Figure 3:**
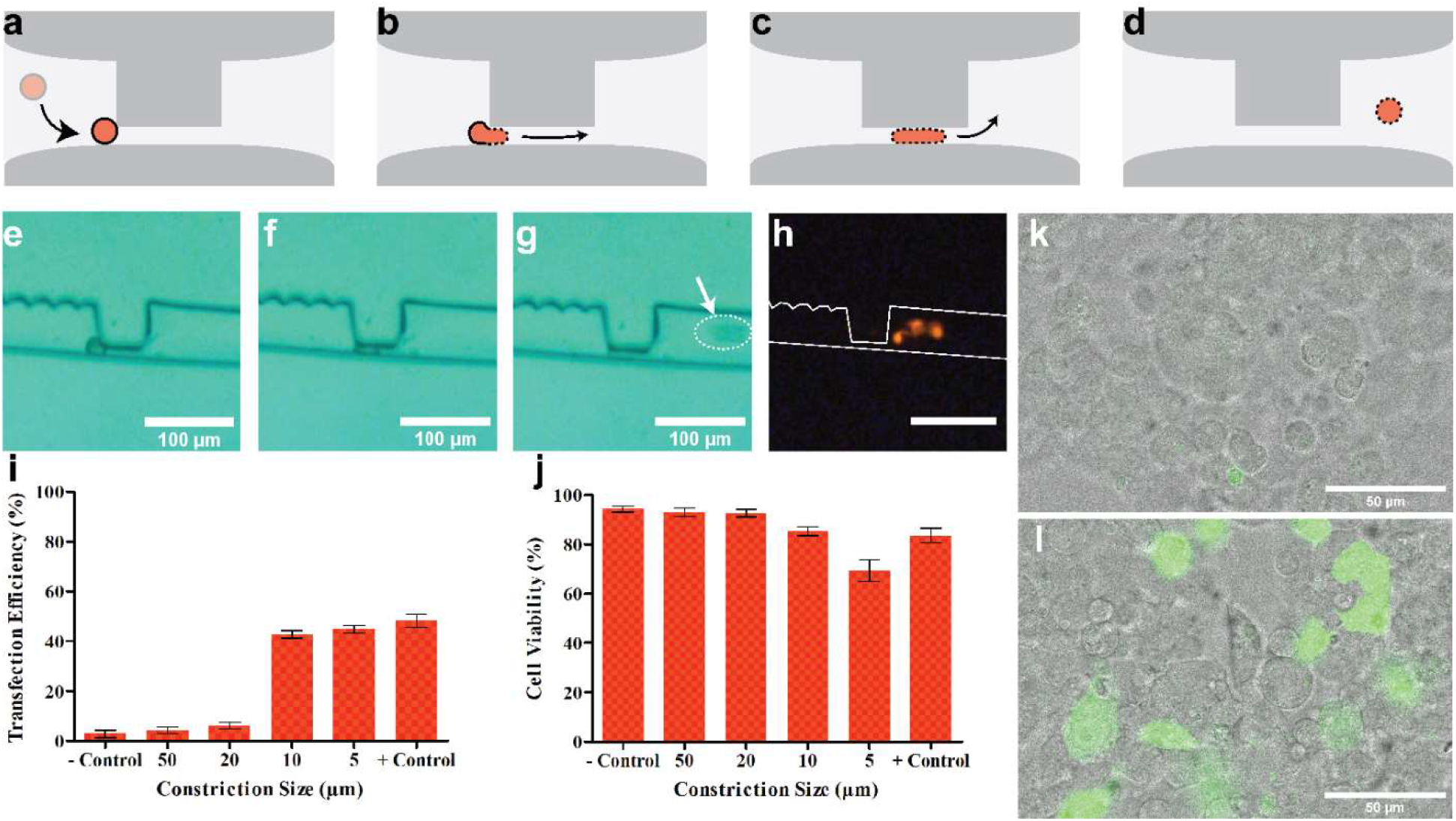
a-d) schematics showing the internal structure of the compression region within a collapsed microfluidic channel with a protruding micro-feature in the shape of a square. A typical cell is represented in orange where the sequence of schematics tracks the movement of the cell under a fixed fluid flow through the temporary constriction formed between the micro-feature and the opposing channel sidewall. The cell wall is shown to become permeable after experiencing rapid deformation. e-f) are a sequence of microscope imagery that shows a cell encountering the deforming constriction, undergoing elongation, and lastly escaping on the other side. h) red fluorescence microscopy image of the same frames in (e-g) where a collection of escaped cells fluoresce in red due to membrane permeation as a result of elongation through the constriction. i) bar graph summarizing the percentage of HEK293 FT cells expressing green fluorescence (transfection efficiency) as a function of passing through different size constrictions. Negative control corresponds to cells that did not enter the channel, and positive control represents cells treated with a commercial transfection reagent. j) bar graph showing cell viability (%) as a function of being flown through different size constrictions. Negative control represents cells not flown through the channel and positive control represents cell viability after treatment with a commercial transfection reagent. k) confocal microscopy bright-field image superimposed on green-fluorescence image of cells from the negative control group of (i) and (j). l) confocal microscopy bright-field image superimposed on green-fluorescence image of cells passed through a 10 µm constriction in the presence of GFP-coding plasmid. Both (k) and (l) were collected after 48 h of incubation under optimal cell proliferation conditions.

### HEK293 FT cell DNA transfection

Once the device operation was successfully verified, the next step was to utilize it to genetically modify the HEK293 FT cell line with a reporter molecule-coding plasmid DNA in order toverify that cell function was not negatively affected. We selected a commercially available GFP-coding plasmid as the target for intracellular delivery.

Cells were resuspended in 4% FBS DMEM medium containing 0.1% Pluronic-F68 and the GFP-coding DNA plasmid. The constriction size was adjusted to 50 (fully open channel), 20, 10, and 5 µm, and in each case, 1 mL of 10^6^ cells/mL was flown through the constriction at 12 µL/min flowrate and collected at the outlet. The collected samples were then incubated for 4 h in a cell incubator, and then transferred through centrifugation to a fully supplemented DMEM solution to remove non-internalized plasmid DNA. The cells were then allowed to incubate in well plates in a cell incubator for 44 h (for a total of 48 h). After incubation, the cells were collected and a portion was imaged using confocal microscopy, while the other portion was stained and analyzed using a flow cytometer (raw data in Figure S5). The results were compiled in Figure 3i-l. We also utilized a commercially available lipid-based cell transfection reagent (EL Transfection Reagent, Abbexa) on a separate culture using the suppliers’ protocol to act as a positive control. Cells that were not injected into the channel but incubated with the DNA plasmid for a similar amount of time (48 h) were used as a negative control. Results in Figure 3i show that as expected, the negative control showed negligible fluorescence in the green channel, while the cells transfected with the commercial reagent showed a 48 ± 4% transfection rate. Cells flown through the 50 and 20 µm showed green fluorescence values similar to the negative control which was expected since no significant deformation occurred to the cells. On the other hand, cells that were flown through the 10 and 5 µm constrictions exhibited green fluorescence values close to the values of the positive control, with the 10 and 5 µm samples showing values of 42 ± 2 and 44 ± 2% respectively. Successful transfection was further confirmed using confocal microscopy where the cells from the 10 µm sample shown in Figure 3k exhibit strong green fluorescence compared to the negative control sample shown in Figure 3l.

In order to test the effects of this transfection method on cell viability, we used Trypan Blue staining assay on the samples after treatment. For cells flown through the channel (for all constriction sizes), the samples were taken and allowed to incubate at room temperature for 30 min to insure complete healing of cell membranes.^[5]^ After that, 10 µL of the sample was taken and mixed with 10 µL of 0.4% Trypan Blue under an optical microscope. Both stained and unstained cells were counted manually and a viability rate (%) was established for each sample as shown in Figure 3j. Similarly, for both positive and negative control, the cells were suspended in solution (after 30 min of exposure to the transfection reagent for the positive control) and then allowed to sit for 30 min at room temperature following which the same procedure was carried out with the Trypan blue stain. Cell viability was calculated as the inverse of the number of blue-stained cells as a percentage of the total number of cells counted. As the figure shows, cells demonstrated less viability as the constriction size decreased. We posit that a larger percentage of the cells that underwent more extreme morphological deformation (i.e. 5 µm constriction) sustained irreversible membrane damage that did not heal over time as compared to larger constriction sizes that did not induce an as extreme morphological deformation as the smallest constriction size. This observation is consistent with^[19]^ where cells that underwent extreme membrane perturbations were less likely to remain viable. We also observed a small decrease in viability for the sample treated with the commercial transfection reagent, which was expected since lipid-based transfection reagents are known to have a slight cytotoxic effect on the treated cells, especially at long exposure times^[30]^. From Figure 3i and j, we surmised that the constriction size that produced the highest transfection rate while maintaining a relatively low rate of cell damage for HEK293 FT cells was the 10 µm constriction size.

### Adherent cell line transfection

Once successful transfection was confirmed on HEK293 FT cells using the described device architecture, two other adherent cell lines were tested similarly using the same DNA plasmid construct. MDA MB 231 and MCF 7 breast cancer cells are some of the most commonly used cell lines in breast cancer research and are readily cultured in DMEM. Since both cell lines are adherent, we included an extra trypsinization step during culture pre-treatment as outlined in the Experimental section of this article. Additionally, we established that the 10 µm constriction size was the most optimal for HEK293 FT transfection in terms of maximizing the rate and maintaining a relatively high cell viability. We decided to use the same constriction size to transfect MDA MB 231 and MCF 7 cell lines even though there are size variations between the different strains. For the positive control, we used the same commercial lipid-based transfection reagent to establish a reference point for comparison. The results of the transfection experiment are displayed in Figure 4. The raw flow cytometry data is shown in Figure S6. Figure 4 also includes the transfection results for HEK293 FT from the previous experiment for comparison. The interesting results were related to the MDA MB 231 cells which showed a marginally larger transfection rate using the device (47 ± 5%) as compared to the commercial transfection reagent (39 ± 5%). It also showed the highest transfection rate using the device as compared to the other tested cell lines. On the other hand, the commercial reagent resulted in a higher transfection rate in MCF 7 cells (56 ± 10%) compared to the device efficiency (44 ± 5%). Overall however, the device resulted in a more or less consistent performance across all the cell lines, with transfection rates in the range of 40-50%. Therefore, it was concluded that the mechanism of transfection is cell species agnostic at least for immortalized cell lines. This is in line with other similar studies showing the successful transfection of multiple species of cell lines via physical cell deformation^[5, 18, 29]^.

**Figure 4:**
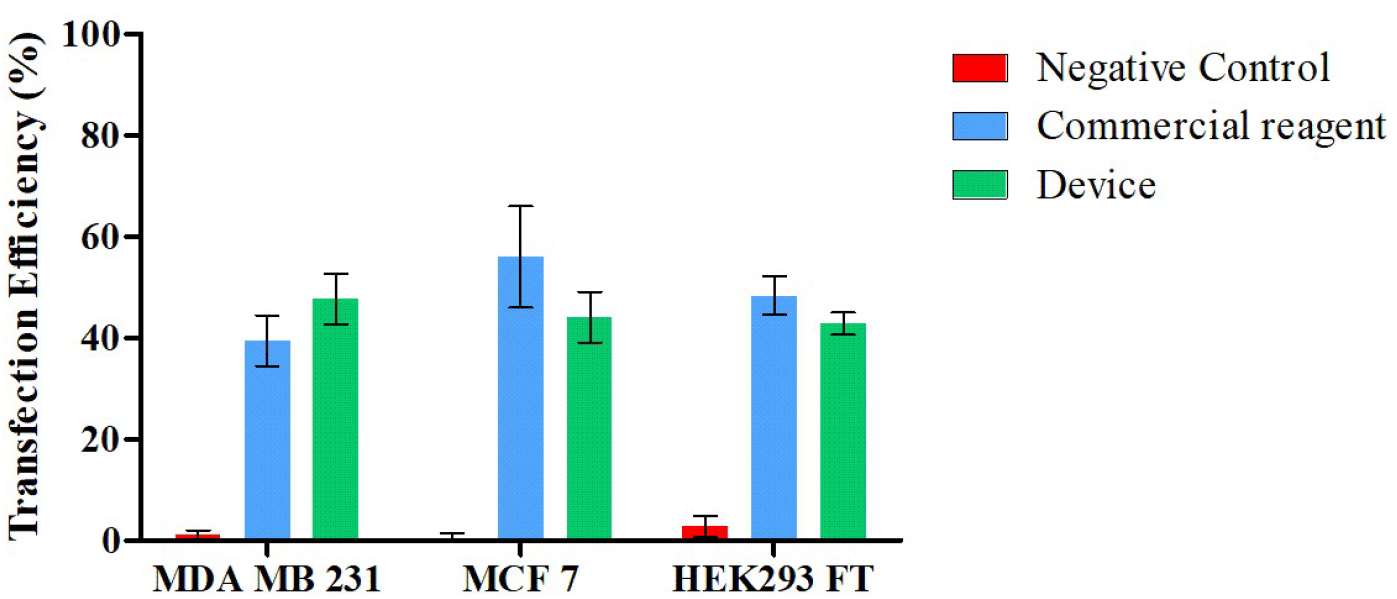
Bar graph representing the GFP plasmid transfection rates as demonstrated by green fluorescence for MDA MB 231, MCF 7, and HEK293 FT. The red bars represent the negative control sample for each where the cells were pre-treated and trypsinized, but not injected into the microfluidic device. The blue bars represent the transfection rate for cells treated with the commercial transfection reagent. Finally, the green bars represent the transfection rates for cells that were flown through the channel with a 10 µm constriction. All experiments were done in triplicates (*n* = 3) as indicated by the error bars.

### Intracellular delivery of fluorescent nanoparticles

What sets physical methods of intracellular delivery (and more specifically, membrane perturbing methods) apart from other biochemical methods is their ability to deliver a variety of materials intracellularly as compared for example to viral or lipid vectors, which are traditionally limited to DNA or RNA payloads. This is important since inorganic macromolecules and nanoparticles serve a range of important functions in cell therapeutics and diagnostics such as fluorescent labelling and tracking, magnetic tagging and control, alteration of cellular function, and in tissue engineering.

Here, we chose commercial 20 nm large gold nanoparticles (AuNPs) tagged with GFP as a model nanoparticle to deliver intracellularly using the physical deformation method. The GFP tag makes it easier to track and quantify the intracellular delivery of the AuNPs and consequently derive the delivery efficiency (%). Pre-treatment of a HEK293 FT culture was performed exactly as outlined in the previous experiments. Instead of loading the carrier DMEM solution with the DNA plasmid, we instead mixed in the AuNPs at a working concentration of 26 µg/mL. Using the device with a constriction size of 10 µm, we passed 1 mL of cells at a cell density of 10^6^ cells/mL. The cell samples that were collected at the outlet were allowed to sit at room temperature for a period of 30 min. The cells were then fixed using a solution of 4% formaldehyde in order to preserve the state of the cells immediately after the intracellular delivery process. The cell samples were then analyzed using flow cytometry. A portion of the fixed cells were also stained using PI and deposited between two glass slides for imaging under a confocal microscope. The results are compiled and shown in Figure 5 while the raw flow cytometry data is shown in Figure S7.

**Figure 5:**
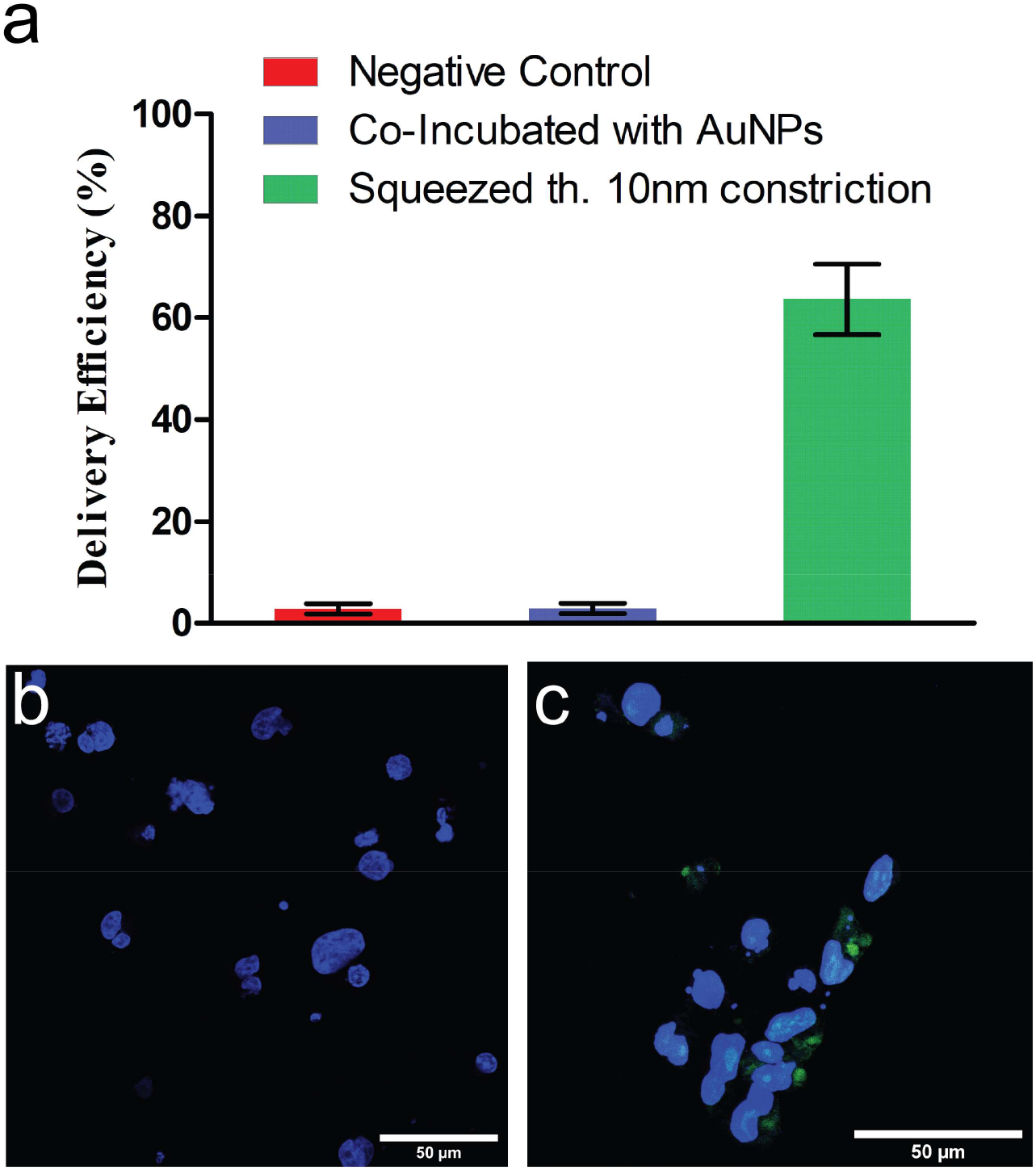
a) is a bar graph representing the green fluorescence intensity which indicated the delivery efficiency of GFP-tagged AuNPs. Negative control (red) shows HEK293 FT cells background fluorescence. Cells co-incubated with AuNPs (blue) were not injected through the channel. Cells squeezed through a 10 µm constriction through the channel (green) were also co-incubated with AuNPs. b) corresponds to cells that were co-incubated with the AuNPs but not squeezed through the channel under a confocal microscope. c) shows cells that were indeed squeezed through the 10 µm constriction under the confocal microscope. Both (b) and (c) have a scale bar of 50 µm length in the bottom right corner of the images. All cells were stained with Hoechst 33342 to highlight the nuclei of cells, while the green color indicated green fluorescence by the GFP-coated AuNPs.

In Figure 5a, the negative control sample was comprised of cells that were not exposed to AuNPs to show background fluorescence. Cells that were co-incubated with the AuNPs but not injected through the channel also showed negligible green fluorescence after washing away the non-internalized AuNPs. These cells are also shown in Figure 5b under confocal microscopy. Cells that were flown through the channel with a 10 µm constriction size showed a relatively high green fluorescence value at around 63 ± 7%. Confocal microscopy images (Figure 5c) confirm the presence of green fluorescing clusters aggregating around or very close to the cell nuclei.

## Conclusions

In this work, we outlined the fabrication of an intracellular delivery microfluidic device that operates on the principle of cell deformation to disrupt the membrane through a constriction in the channel. The entirely elastic composition of the device allowed for the creation of temporary channel constrictions on demand with the desired size using mechanical compression of the device in certain regions of the channel. By utilizing the device, we demonstrated the successful transfection of three mammalian cell lines with a plasmid construct coding for GFP. Transfection efficiencies were achieved that were close to, or even surpassing the performance of a common commercial transfection reagent while maintaining high cell viability throughout the duration of the experiments. We also demonstrated that the device was able to deliver inorganic metal nanoparticles (AuNPs) intracellularly to HEK293 FT cell lines unlike the commercial transfection reagent. By controlling the constriction diameter and chemically passivating the inner walls of the channel, we were able to clear the vast majority of clogs and prevent cell and debris agglomeration, allowing the device to be used multiple times for multiple cell lines after proper decontamination. This device is characterized by a very small footprint and minimalistic operational requirements (comprised of a power source and circuitry to drive the tiny linear actuators). It is also highly flexible and durable, allowing it to be easily integrated in most lab-on-a-chip assemblies for cell therapy. The utility of this dynamic constriction forming mechanism will not be limited to mechanical intracellular delivery. The real-time application of mechanical force can be used to study the effect of mechanical forces on cells, or act as micro-tweezers that hold selected cells in place while measurements are carried out, or for operations such as optoporation, etc. We believe that the application of mechanical force on key regions of microfluidic assemblies will generate many novel and exciting applications for the field of microfluidics in the future.

## Experimental

### Device fabrication and mechanical actuation

First, the PDMS channel geometry and micro-features were designed using LayoutEditor software package. Inlet and outlet were defined as 5x5 mm squares and connected through a 100 µm wide, and 39 mm long channel. The micro-features were defined as two polygon-shaped protrusions with the first having dimensions of 30x50 µm and the second having dimensions of 50x50 µm. The two polygons were separated by a 100 µm region of triangular spikes with a base 20 µm long and a height of 20 µm. The mask was copied onto a chrome-coated photolithography mask using a mask writer (Heidelberg, DWL66FS). A flat silicon wafer (p-type 4”, Siegert) was coated with a 100 µm thick layer of SU-8 and a mask aligner (EVG 620) was used to expose the pattern on the mask at 600 mJ/cm^2^. The SU-8 was then hard baked at 65°C for 15 min, and then developed using an SU-8 developer solvent. The wafer was then cleaned using isopropanol and water and then dried using a N_2_ stream.

PDMS (SYLGARD 184 elastomer kit, Dow Chemical) was mixed at a ratio of 15:1 elastomer:cross-linker to a total of 8 mL and poured carefully over the SU-8 pattern. A similar mixture was prepared and poured over another flat silicon wafer to compose the second device layer. Both mixtures were degassed in a vacuum chamber and then allowed to cure on a hotplate at 95°C for 40 min. Once the PDMS slabs solidified, they were carefully peeled off the wafers, and the slab containing the channel engraving was punctured with a 1.25 mm puncher at the center of the inlet/outlet reservoirs. Both slabs were then exposed to O_2_ plasma (300 W, 20 sccm O_2_, 5 sccm N_2_) for 15 min to fully oxidize. Both PDMS slabs were then manually bonded to each other and degassed in a vacuum chamber to remove any trapped bubbles. The bonded slabs were cured again over a hotplate at 95°C for 40 min to increase bonding strength. Finally, a CNC-cut dog-bone shaped stencil was used to cut out the complete channel from the slabs. Teflon tubing was attached to each punch-hole at the inlet and outlet to provide fluid access to the channel.

Acrylic frame and base were cut from a 5 mm thick transparent acrylic sheet using a CNC machine and then assembled as shown in Figure 1b. The PDMS device was placed in the middle of the device and capped with another clear acrylic piece to prevent any bulging in the z-axis (perpendicularly up). Two miniaturized linear actuators (Hybrid bipolar 5V DC, 12.5 mm stroke, 8.7 µm step-size, 10 N holding force @ 4V, dimensions: 22 mm x 16.5 mm x 4.5 mm) were attached to the acrylic frame and connected to two acrylic pusher arms (Figure 1b and d). The motors were driven by an X-Nucleo-IHM06A1 controller board and an Arduino Uno microcontroller using custom-written code.

### Channel chemical passivation

For all fluid manipulations, a Fluigent pneumatic microfluidic control system (MFCS-EZ) was used. A solution of 5:1:1 H_2_O:HCl:H_2_O_2_ was flown through the channel at a flowrate of 10 µL/min for 30 min, followed by DI water for 30 min. Successively more concentrated ethanol solutions in DI water (50%, 70%, 90%, 100%) were flown through the channel for 10 min each at the same flowrate, with the last solution (100%) being flown for 30 min to completely dehydrate the channel. A solution of 4% 1H, 1H, 2H, 2H-Perfluorooctyltriethoxysilane (Sigma) in ethanol was flown for 30 min to silanize the interior of the channel. This was followed by a 30 min long rinse using 100% ethanol and another subsequent rinse using DI water. The device was then ready for cell experiments.

### General cell culture conditions

All cell lines (HEK293 FT, MDA MB 231, MCF 7) were procured from frozen samples (−80°C) and thawed in a warm water bath (37°C). The cells were transferred to high glucose Dulbecco’s Modified Eagles Medium (DMEM) supplemented with 20% fetal bovine serum (FBS), insulin, and essential amino acids with penicillin/streptomycin after removing the cryogenic media using centrifugation at 1500 rpm for 5 min. The cells were cultured in 145/20 mm cell culture plates until confluency reached ∼80% through visual inspection.

### Cell culture pre-treatment protocol

Before initiating experiments that involved the use of the microfluidic device, the following cell culture pre-treatment protocol was followed to reduce cell aggregation. Cell cultures were extracted after reaching 80% confluency, and then immersed in a solution of 1 mg/mL of DNase I (Abbexa abx082222) in order to digest any extracellular DNA material and reduce intercellular adhesion. This was followed by a washing step in 1x PBS and a second immersion step in 0.02% Ethylenediaminetetraacetic acid (EDTA, Sigma) to chelate Ca^+^ ions (that cause cell aggregation) and to deactivate any remaining DNase I. Two more washes in 1x PBS were performed, followed by cell resuspension/trypsinization as required.

### Trypsinization and cell resuspension

For cells that needed trypsinization, Biowest Trypsin-EDTA 10x was diluted 1:10 in 1x PBS and used to trypsinize the cells for 2 min, after which the cells were pelleted and transferred to fresh DMEM media for further processing. HEK293 FT cells were simply agitated via pipetting to resuspend the cells for further processing.

### General device usage

For all experiments that involved the use of the PDMS device, the following general protocol was followed unless otherwise stated in the text. Initially, the device constriction was set to the desired value using the linear actuators and control circuitry. After that, cells were detached from the culture plate and transferred into DMEM supplemented with 4% FBS and 0.1% Pluronic-F68 (to help repair membrane damage and reduce intercellular adhesion). The cells were counted to insure they are in the range of 10^6^ cells/mL. The desired cargo (plasmid or AuNP) was added to the intracellular delivery medium at this stage and 1 mL of the medium was injected into the device channel using the Fluigent system at a flowrate of 12 µL/min. If clogging was observed at any stage, the device constriction was opened to allow the clogging mass to flow, and then it was reset to its initial constriction size. The output sample was then collected and processed further in accordance with the cargo used.

### DNA plasmid transfection

All experiments involving the transfection of GFP plasmid utilized Altogen Biosystems GFP-expressing plasmid DNA (catalogue No. 4060). For each 1 mL of sample containing ∼10^6^ cells/mL, a concentration of 0.8 µg/mL of plasmid was added to the transfection medium. For all positive-control experiments, a similar amount of DNA plasmid (0.8 µg) was mixed at a ratio of 1:2 w/v% with the commercial transfection reagent (EL Transfection Reagent, Abbexa abx098880) and the protocol provided by the supplier was followed without major deviations.

Cell volumes treated with the PDMS device for all constriction sizes were collected and then allowed to rest at room temperature for 30 min. 10 µL of each sample was taken and mixed with 10 µL of Trypan Blue stain on a microscope slide, after which it was viewed under an inverted microscope. Blue-stained cells were counted as a function of the total number of cells, and the viability was reported as such. The remaining cell volume was placed in a 6 well plate and incubated under cell culture conditions for 3.5 h. This was followed by draining the transfection medium and adding fresh fully supplemented DMEM, and the cells were allowed to further incubate at cell culture conditions for 44 h (for a total of 48 h). The cells were then analyzed using a flow cytometry system (BD Accuri C6 Plus). Flow cytometry data was analyzed and plotted using Python and associated libraries.

### AuNPs intracellular delivery

All experiments involving the intracellular delivery of fluorescent AuNPs utilized Nanopartz 20 nm functionalized fluorescent spherical gold nanoparticles (CF11-20-GFP-FM-50-1) coated with polymer and GFP, with excitation/emission wavelengths of 488/510. Each application involved the use of 26 µg/mL for each sample. After cell treatment with the device, the cells were collected at the output and allowed to sit at room temperature for 30 min. The cells were then washed to remove non-internalized AuNPs and then fixed in a solution of 2% glutaraldehyde in PBS for 10 min at room temperature. After fixation, the cells were analyzed for fluorescence using flow cytometry.

## Author Contributions

**Hashim Alhmoud:** Conceptualization, methodology, investigation, writing – original draft, funding acquisition **Mohammed Alkhaled:** Conceptualization, formal analysis, visualization, writing – review & editing **Batuhan E. Kaynak:** Software, formal analysis, visualization, writing – review & editing **M. Selim Hanay:** Resources, writing – review & editing, supervision, funding acquisition.

## Acknowledgements

The authors would like to acknowledge project funding by the Marie-Curie Actions Horizon 2020 and the Scientific and Technological Research Council of Turkey (TÜBİTAK) under the joint funding scheme titled “Brain Co-circulation Scheme 2 (2236-CoFund)”, project number **120C056**. This project has also received funding from the European Research Council (ERC) under the European Union’s Horizon 2020 research and innovation programme (grant agreement **n° 758769**). The authors would also like to acknowledge the roles of Mr. Ege Erdem and Mr. Abdulrzak Masrani for their help in building the electronic system driving the linear actuators.

