## Supplementary Information for "Leveraging the Elastic Deformability of Polydimethylsiloxane Microfluidic Channels for Efficient Intracellular Delivery"

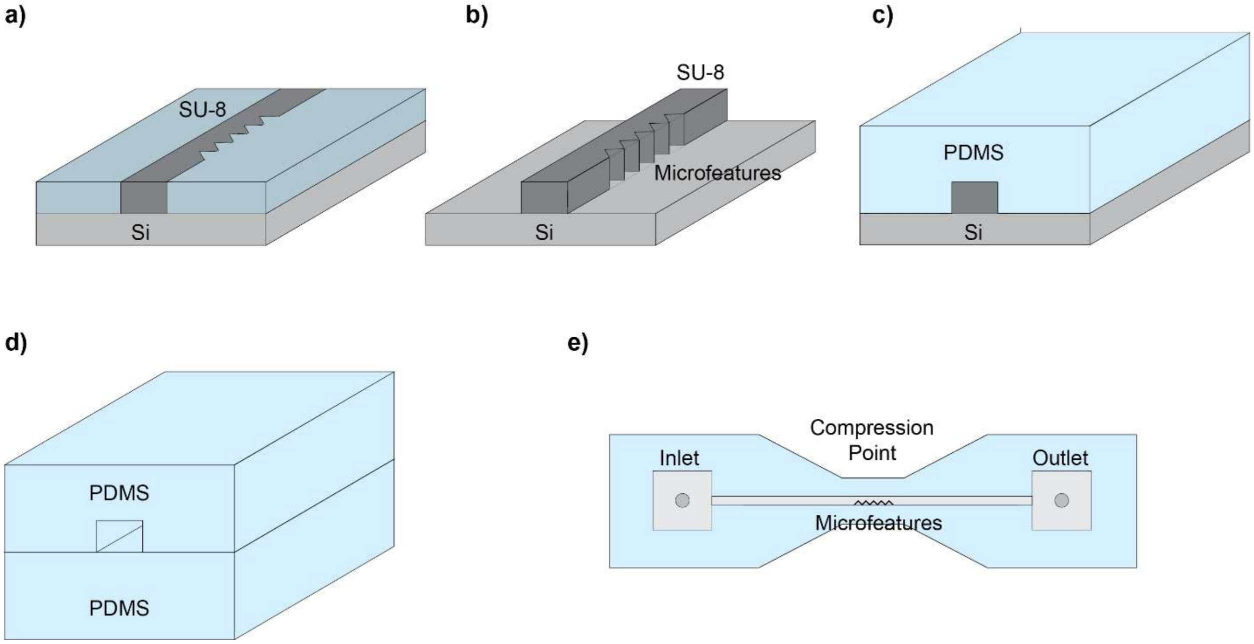


*Figure S1: Schematic diagram showing the fabrication of the PDMS device. a) A 100 µm layer of SU-8 photoresist is spun-coated uniformly on a flat silicon wafer. The SU-8 layer is exposed to UV light through a photolithography mask to define the negative of the microfluidic channel with a 100 µm width. The micro-features are defined within the compression region on the photolithography mask at this stage. b) The wafer is hard-baked at 65ºC for 15 min and the unexposed SU-8 is dissolved by immersion into SU-8 developer for 10-15 min. c) A mixture of 15:1 PDMS:cross-linker is uniformly coated to the top of the wafer to a thickness of 1 mm. The PDMS is then degassed in a vaccum chamber and then allowed to cure at 95ºC for 40 min on a hotplate. Once the PDMS is cured, it is carefully pealed off the silicon wafer. This forms the top layer of the device and includes the engraving of the channel defined by the cross-linked SU-8. At this stage 1.25 mm holes are punched through at either end of the channel to form the inlet and outlet. d) Another completely flat 1 mm PDMS layer using the same 15:1 ratio is prepared to form the bottom layer of the device. Both layers of the device are oxidized in $O_2$ plasma and then bonded together under vaccum, and then cured at 95ºC for 40 min. e) Once the device is completely bonded, it is tested for leaks and then cut out into a dog-bone shape using a stensil ensuring that the micro-features occupy the middle-point of the narrow region of the dog-bone to form the compression region.*

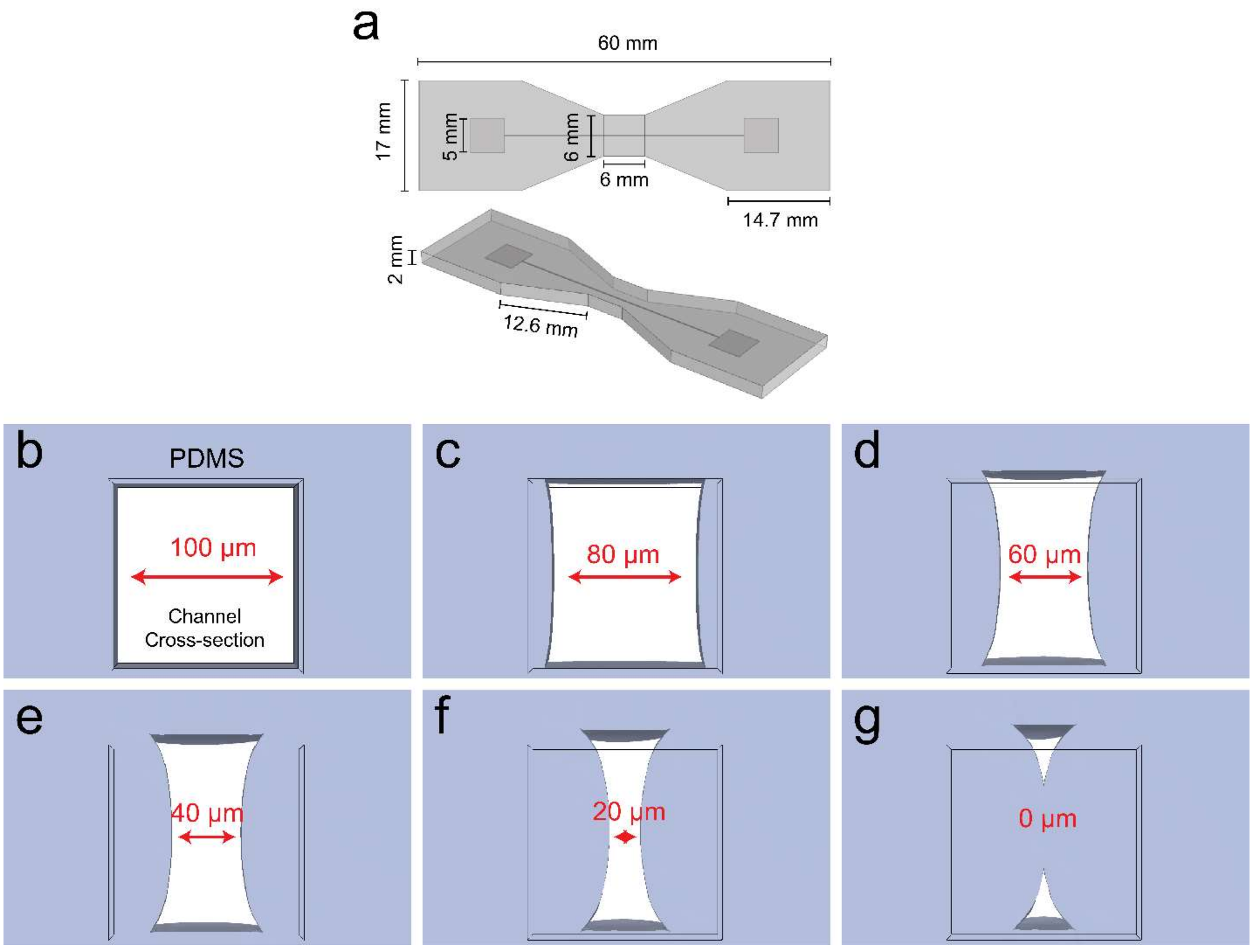


*Figure S2: Panel (a) schematic describing the exact dimensions of the PDMS device used in the experiments. Panels (b-g) is a rendered cross-sectional view of the channel at the center of the compression region obtained through simulation. The successive panels show the bulging profile of the channel sidewalls as the external mechanical force is increased. This is a byproduct of the top and bottom of the sidewalls being geometrically fixed to the ceiling and floor of the microfluidic channel, and with any vertical translation being blocked. Even with the channel fully closed, there remains two open portions at the top and bottom (g) that will sustain a small amount of fluid flow (i.e. even when the channel is fully closed, we observe a small flowrate < 1 µL/min flowing through the channel outlet).*

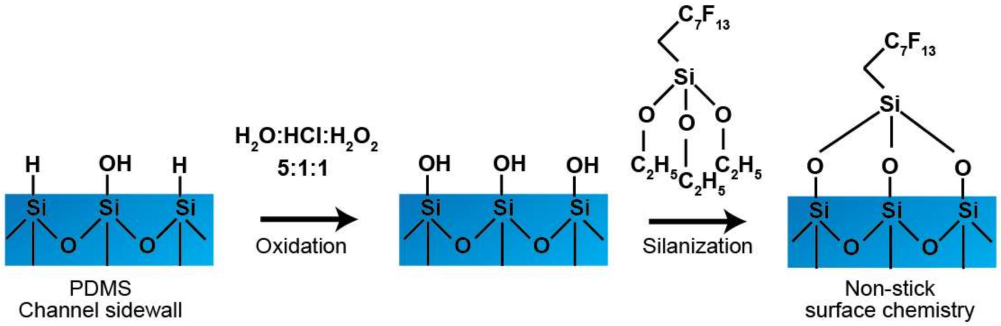


*Figure S3: Schematic depicting the reaction scheme used to passivate the surface of the inner walls of the microfluidic channel. The PDMS surface is usually covered with silicon hydride and silicon hydroxide groups. Exposing it to an oxidizing solution of $H_2O$:HCl:$H_2O_2$ (5:1:1) will cause the entire surface to become oxidized and silicon hydroxide groups become dominant. The surface is then dried and exposed to 1H, 1H, 2H, 2H-Perfluorooctyltriethoxysilane (4% v/v% in ethanol) for 30 min at room temperature. The reaction lead to the formation of Si-O-Si groups thus covalently bonding the fluorosilane to the surface of the inner walls of the channel. The long fluorocarbon chain (-$C_7F_{13}$) effectively passivates the surface and prevents ionic and Van der Waals interactions with the surface.*

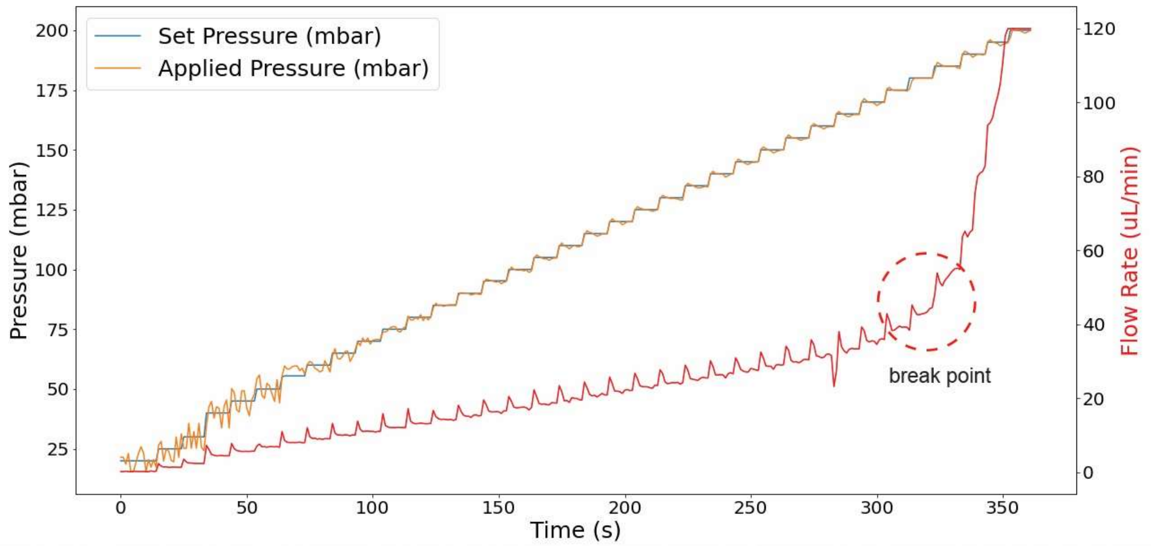


*Figure S4: Plot that shows the maximum flowrate tolerated by the PDMS device before developing a leak. The first y-axis describes the pressure (in mbar) applied to a fluid reservoir to push it through the PDMS microfluidic channel (Fluigent) where the set pressure is defined using the controller software, and the applied pressure is what is actually outputted by the air pump. The second y-axis is the output of a flowrate sensor attached between the fluid reservoir and the microfluidic channel. The salient feature of this plot is a clear "break point" where a leak develops between 75 -100 mbar. This is indicated by a sharp increase in the flowrate after 325 s which correlates with a maximum flowrate of ~ 42 µL/min.*

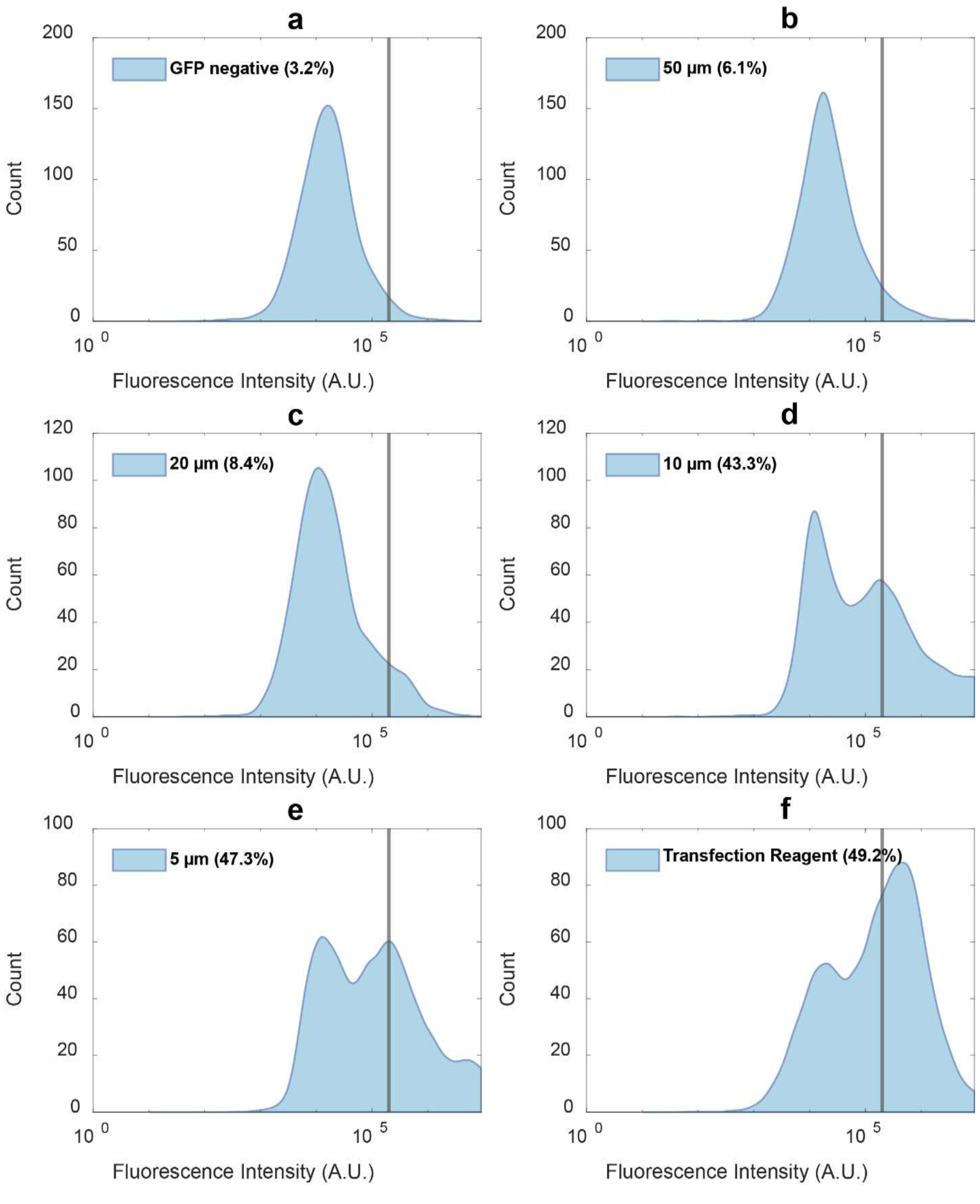


*Figure S5: Flow cytometry results of fluorescence in the green channel for HEK293 FT cells treated using the device in the presence of the GFP-coding plasmid. Panel (a) shows the fluorescence of cells not treated with the device to establish a threshold for HEK293 FT background fluorescence, which is indicated by the vertical red line positioned at $2 \times 10^5$ A.U. Panels (b-e) represent cells that passed through the PDMS channel at different constriction sizes (50, 20, 10, and 5 μm). Cells that fluoresced with an intensity larger than the established threshold were counted as successfully transfected. The red percentage value in each panel represents the amount of cells with fluorescence above background. Panel (f) shows GFP fluorescence for HEK293 FT cells transfected using a commercial reagent as a positive control.*

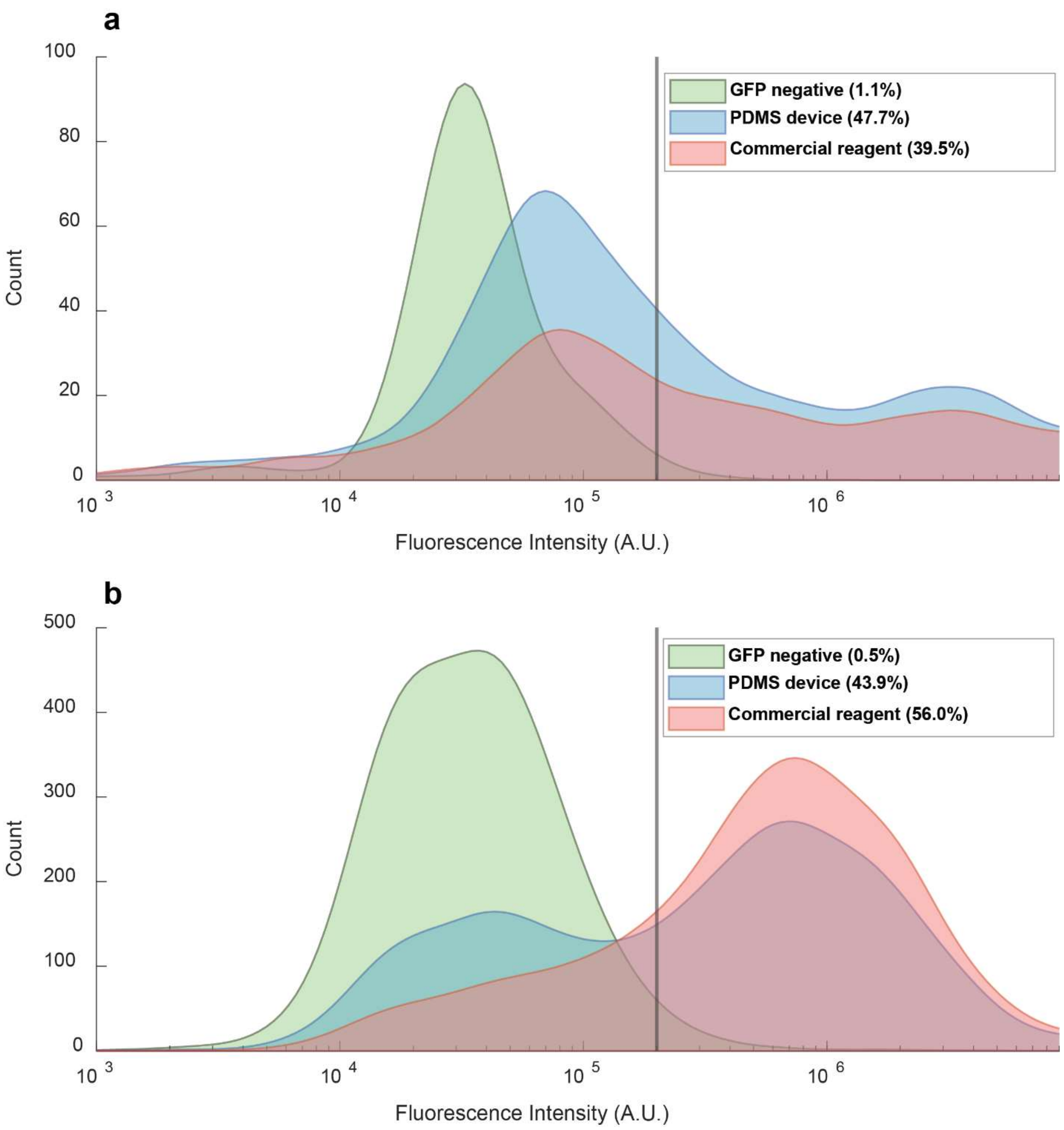


*Figure S6: Flow cytometry results of fluorescence in the green channel for (a) MDA MB 231 cells and (b) MCF 7 treated using the device (10 µm constriction size) in the presence of the GFP-coding plasmid. The results are compared to a positive control which is a commercial lipid-based transfection reagent.*

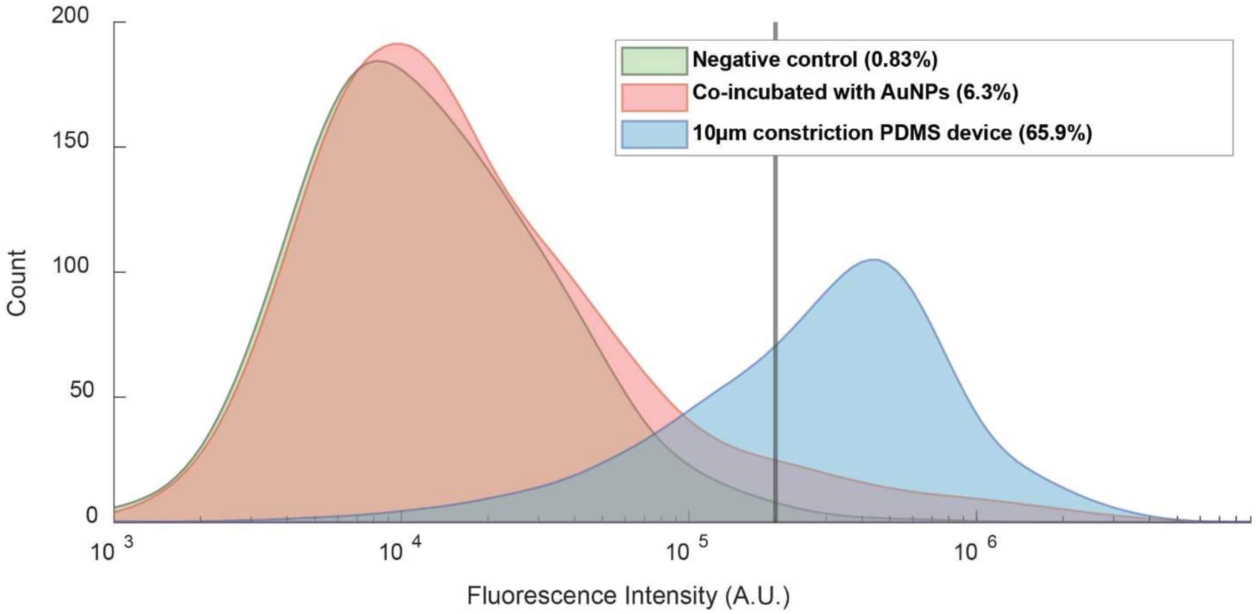


*Figure S7: Flow cytometry results of green channel fluorescence for HEK293 FT cells treated with the PDMS device in the presence of GFP-labeled 20 nm AuNPs. Negative control labeled sample (blue) are untreated cells used to establish threshold for significant fluorescence. The co-incubated sample (orange) refers to cells co-incubated with AuNPs but not injected through the device channel. The sample squeezed through 10 µm constriction (green) refers to cells co-incubated with AuNPs and injected into the device channel with a 10 µm constriction.*
